# Generation of densely labeled oligonucleotides for the detection of small genomic elements

**DOI:** 10.1101/2024.03.15.583980

**Authors:** Clemens Steinek, Miguel Guirao Ortiz, Gabriela Stumberger, Annika J. Tölke, David Hörl, Thomas Carell, Hartmann Harz, Heinrich Leonhardt

## Abstract

The genome contains numerous regulatory elements that may undergo complex interactions and contribute to the establishment, maintenance, and change of cellular identity. Three-dimensional genome organization can be explored with fluorescence in situ hybridization (FISH) at the single-cell level, but the detection of small genomic loci remains challenging. Here, we provide a rapid and simple protocol for the generation of bright FISH probes suited for the detection of small genomic elements. We systematically optimized probe design and synthesis, screened polymerases for their ability to incorporate dye-labeled nucleotides and streamlined purification conditions to yield nanoscopy-compatible oligonucleotides with dyes in variable arrays (NOVA-probes). With these probes, we detect genomic loci ranging from genome-wide repetitive regions down to non-repetitive loci below the kilobase scale. In conclusion, we introduce a simple workflow to generate densely labeled oligonucleotide pools that facilitate detection and nanoscopic measurements of small genomic elements in single cells.

## INTRODUCTION

In recent years, multiple layers of mammalian genome organization ranging from preferential positions of chromosomes in the nucleus to active and inactive compartments and small-scale interactions between individual loci have been uncovered ^1–6^. An intricate interplay of chromosome territories, topologically associated domains and regulatory elements defines cellular identity in development and disease ^7–11^. While current methodologies reliably probe pairwise and multi-contact DNA-DNA interactions, deciphering complex 3D chromatin organization in single cells remains challenging, particularly in the kilobase range ^12–14^. Thus, there is a growing demand for increased sensitivity to detect and study DNA elements in the 3D context of individual nuclei.

The state-of-the-art for mapping chromatin contacts are chromatin capture assays ^15–18^. These methods are especially powerful as they detect contacts within large-scale genomic regions with a resolution ranging from one kilobase down to the nucleosome level but typically rely on population averages ^19–21^. However, early efforts to probe chromatin contacts in single cells using chromatin capture assays have revealed extensive cell-to-cell variations within the same population ^22^. Inter-cell variation of 3D chromatin structures has been observed in multiple imaging studies, which is consistent with the transient nature of chromatin contacts revealed by live-cell imaging ^23–31^. Therefore, chromatin capture assays need to be complemented with sensitive imaging methods to comprehensively address the dynamics and function of chromatin conformations.

Since their advent, microscopy and fluorescence in situ hybridization (FISH) have shed light on the spatial distribution of chromatin in single cells and identified chromosomal abnormalities in malignant cells and tissues ^32–35^. Although fluorescence microscopy has facilitated studies on large-scale chromatin structures, the detection and resolution of small regulatory elements with traditional FISH methods remains challenging ^23,36^. In past works, FISH probes have often been generated from bacterial artificial chromosomes (BACs) or yeast artificial chromosomes (YACs) using polymerases in random priming or nick translation reactions ^37–42^. However, the size of BAC or YAC probes limits the genomic resolution and is, therefore, not suitable for the detection of short regulatory DNA sequences ^43^. Recent advances in synthetic DNA production and the availability of whole genome datasets have ushered in a new era of oligonucleotide-based FISH methodologies ^44–46^. Variations of oligonucleotide-based FISH (oligoFISH) utilize barcoded primary pools and fluorescent secondary readout probes to sequentially detect genomic loci ^30,31,46,47^. Although this approach has enabled considerable advancements in understanding chromatin architecture, the usage of single-labeled secondary probes limits the detectable target size and spatial resolution. Signal amplification has been achieved through the hybridization of multiple secondary probes to a prolonged primary strand (SABER-FISH) but has mostly been used for the detection of RNA ^48^. We hypothesized that the direct coupling of multiple fluorophores to primary oligonucleotides in combination with the elimination of secondary hybridization steps improves the signal-to-noise ratio at DNA loci of interest.

Here, we introduce a protocol to generate nanoscopy-compatible oligonucleotides with dyes in variable arrays (NOVA-probes). Multiple fluorophores are attached to oligonucleotides in a one-step biochemical reaction, thereby considerably shortening the time required for probe generation. The protocol has further been optimized to allow precise control of the labeling density and does not require demanding amplification or purification steps. We applied our probes to detect a variety of genomic loci ranging from large-scale repetitive regions to sub-kilobase single loci using FISH (NOVA-FISH). Compared to previous methods, NOVA-FISH probes can efficiently be produced and allow free choice of fluorophores and flexible adjustment of labeling density to optimize signal detection in super-resolution microscopy.

## RESULTS

### Design and Synthesis of NOVA-Probes

Oligo-based FISH methods have proven valuable in visualizing genomic regions, but the necessity of multiple hybridization steps and/or the use of expensive, end-labeled probes limit their widespread application in nanoscopy. We reasoned that densely labeled oligonucleotide probe sets could be generated with an enzymatic approach in an efficient and cost-effective manner (Figure 1A). To this goal, we hybridized 5’-phosphate-labeled template strands with short primers followed by primer extension and lambda exonuclease-mediated template degradation. Compared with barcoded oligonucleotides and end-labeled probes, our densely labeled oligonucleotides (NOVA-probes) significantly improve signal strength and detectability (Figure 1B-C, and S1).

As our approach depends on the enzymatic incorporation of modified nucleotides into short primers, we compared commonly available DNA polymerases. We measured the incorporation of different dye-labeled nucleotides during extension using commonly available family A (Klenow exo-, Taq) and family B (Q5, Phusion, Therminator) DNA polymerases. Photometric measurements of synthetised probes showed that the highest labeling rates were obtained for all tested modified nucleotides with Therminator DNA polymerase (Figure 1D, Figure S2, S3A-B).

Therminator DNA polymerase is a DNA polymerase that has been derived from the euryarchaeon Thermococcus sp. 9°N and carries mutations in its exonuclease domain (D141A, E143A) and finger domain (A485L) ^49^ (Figure 1E). As a result of these modifications, Therminator DNA polymerase exhibits a decreased discrimination for modified nucleotides and has been used to synthesize a variety of unnatural nucleic acids ^50–53^. To investigate the molecular basis for the observed variations in incorporation efficiencies among our candidates we modeled dye-labeled nucleotides in different conformations in conjunction with finger domains of family A and B polymerases (Figure S4). We noted possible steric clashes between dye-labeled nucleotides and finger domains of family A members, whereas no such clashes were observed with family B polymerases. Using Therminator DNA polymerase, we determined that probes are robustly generated within an hour (Figure S3C). In addition, our approach allows free choice of fluorophore and flexible adjustment of labeling density (Figure S3D).

FISH probes require a high degree of purity since complementary or unlabeled strands will compete with the labeled probe during hybridization and, thus, reduce signal intensity. To remove unbound primers, free nucleotides, and enzymes, we adapted the buffer conditions to selectively yield double-stranded oligonucleotides after extension (Figure S5A-B). Also unlabeled template DNA might block the synthesized probes and thereby prevent their hybridization with the locus of interest. Therefore, we have introduced phosphate groups at the 5’-ends of template strands to mark them for lambda exonuclease-mediated degradation (Figure S5D). Using this approach, template DNA was effectively degraded within 30 minutes (Figure S5E). We then used ethanol-based purification to obtain the single-stranded probe (Figure S5A, Figure S5C).

After establishing a robust workflow, we assessed the number of incorporated fluorophores in NOVA-probes. High-performance liquid chromatography (HPLC) analysis revealed that using a low ratio of modified to unmodified nucleotide (25%) in the synthesis reaction yields distinct elution peaks corresponding to the incorporation of increasing numbers of fluorophores (Figure S6).

**Figure 1.**
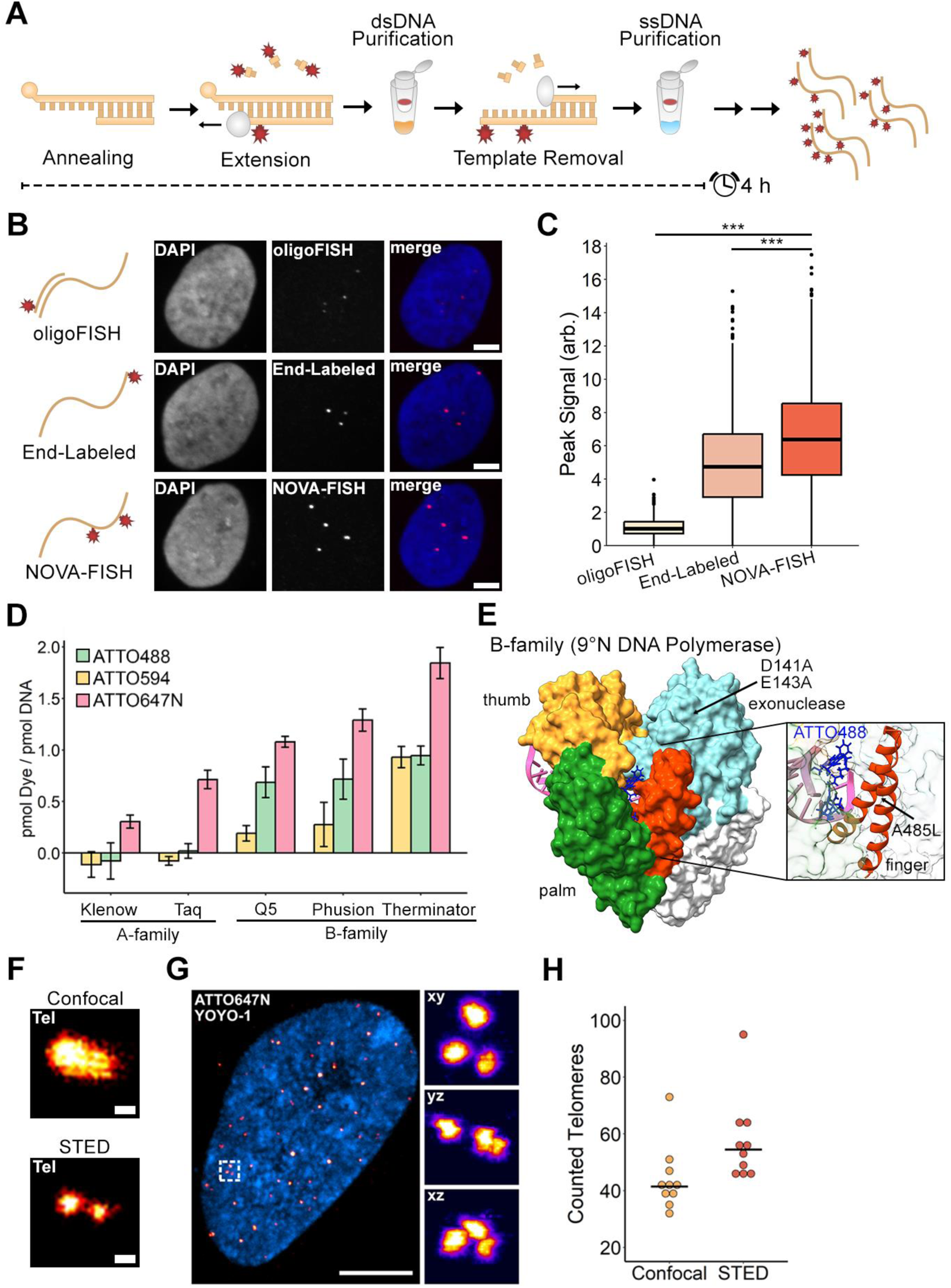
Generating oligonucleotides that carry multiple fluorophores. (A) Schematic workflow of the protocol. Primers are annealed to 5’-phosphorylated template strands and dye-labeled nucleotides are incorporated in a one-step extension reaction. Template strands are then enzymatically removed and the product is purified. (B) Comparing three FISH strategies to tag genomic loci. While oligoFISH uses labeled readout strands for detection, end-labeled and NOVA-FISH probes carry fluorophores in their primary sequences. FISH was conducted in subtelomeric regions in U2OS cells. Scale bars, 5 μm. (C) NOVA-FISH yields bright FISH signals. Related to (B). Number of detected signals: oligoFISH (n = 852), End-labeled (n = 839), NOVA-FISH (n = 1182). Data sets were tested for significance using the Wilcoxon rank sum test with Bonferroni’s correction for multiple testing (*** = p < 0.001). (D) Screening substrate preferences of selected DNA polymerases. Polymerases (Klenow exo-, Taq, Q5, Phusion, Therminator) incorporated dCTP-ATTO488, dCTP-ATTO594, or dCTP-ATTO647N into oligonucleotides using a one to four molar ratio of dye-labeled to unlabeled nucleotides. (E) Crystal structure of the 9°N DNA polymerase in complex with DNA and dCTP-ATTO488. The protein is shown in white with highlighted palm (green), thumb (yellow), finger (orange), and exonuclease (cyan) domain. dCTP-ATTO488 was superimposed on the incorporated nucleotide in the complex. The magnified image depicts dCTP-ATTO488 in the binding pocket. The figure was generated with UCSF Chimera (v.1.17.3, RRID:SCR_015872) accessing 5OMV ^58,59^. (F) Super-resolution microscopy uncovers clustered telomeres. Representative image of two clustered telomeres using confocal microscopy or STED microscopy. Scale bars, 200 nm. (G) Telomere clustering is a common phenomenon in mitotic cells. Representative image of telomeres in IMR-90 cells using 3D-STED microscopy. Detailed view (white box) in three dimensions. Scale bar, 5 µm. (H) Counting telomeres using confocal microscopy or 3D-STED. Related to G. Telomeres were counted in 10 individual cells from three experiments (n = 3). The black lines depict the mean.

### Visualizing telomere clustering below the diffraction limit

Next, we sought to utilize the brightness of NOVA-probes to visualize telomeres below the diffraction limit. We tagged telomeres with telomere-specific NOVA-probes and acquired images using confocal or STED microscopy (Figure 1F). We observed clustered telomeres using STED microscopy, which appear as single entities in confocal images. We then applied 3D STED microscopy to gain further insights into the degree of telomere clustering (Figure 1G). Telomeres in the same cells exhibited considerable heterogeneity in their size and clusters containing multiple telomeres were observed, consistent with previous works ^54–57^. Next, we analyzed the number of detectable telomeres using confocal or STED microscopy (Figure 1H). In comparison to confocal images, STED microscopy detected on average 1.31 times more telomeres (±SD = 0.21), corresponding to clustered telomeres that are only resolved with super-resolution microscopy. Hence, the brightness of NOVA-probes supports demanding super-resolution microscopy to visualize nuanced details of genomic loci with high optical resolution.

### Dense labeling does not affect hybridization efficiency but reduces signal strength

As our workflow yields densely labeled probes, we next tested how the presence of multiple dyes in the probe affects hybridization efficiency. To address this, we generated barcoded probes with increasing labeling densities (Figure 2A-C, Figure S2D). These probes contain dye-labeled sequences that bind to the genome and unlabeled barcodes that hybridize with secondary probes carrying another dye. Using this approach, we can evaluate the brightness of the NOVA-probe signal (green) and the relative number of probes localized at the target region (red) (Figure 2D-E). We found that increasing the number of dye-labeled nucleotides in the probe did not affect the number of bound probes at the locus of interest, as no notable drop in red signal was observed (Figure 2F). However, the brightness of our probes decreases at high labeling densities (Figure 2G). Consequently, densely labeled probes still bind to the region of interest, but short intermolecular distances between fluorophores likely impede signal strength (Figure 2C).

### Establishing densely labeled probes with regularly spaced fluorophores

Our previous strategy yields labeled oligonucleotides in an efficient and cost-effective manner but depends on the occurrence of cytosines in the synthesized sequence. Therefore, we modified our workflow to generate extended probes (xNOVA-probes) that carry fluorophores in a protruding sequence that does not bind to the genome (Figure 3A) ^46,47^. In this design, fluorophores are regularly spaced in the invariable sequence to avoid distance-dependent effects that might diminish the specific brightness. We synthesized probes that either carried one (1x1C), two (2x1C), or three (3x1C) fluorophores and measured their fluorescence signals at the locus of interest (Figure 3B-C). The addition of longer sequences (2x1C, 3x1C) resulted in stronger signals (Figure 3D). With this approach, we observed a steady increase in signal strength at higher labeling densities arguing against short distance-dependent, negative effects in 3x1C sequences (Figure S7).

**Figure 2.**
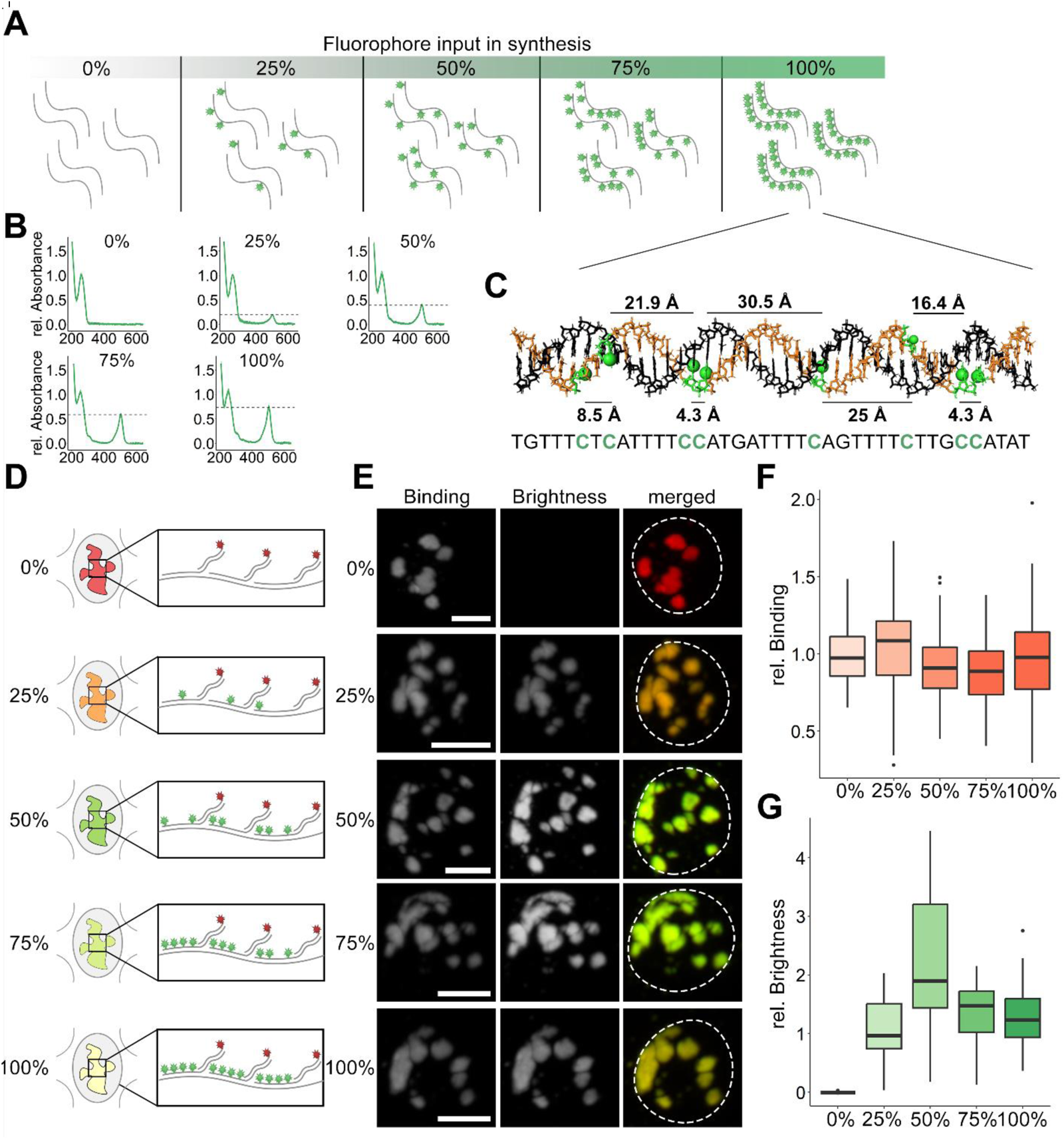
Binding efficiency and brightness of densely labeled probes. (A) Modulating labeling densities during NOVA-probe synthesis. The labeling density is controlled through the ratio of labeled to unlabeled nucleotide (0%, 25%, 50%, 75%, 100%) in the synthesis reaction. (B) Absorption spectra of probes with increasing labeling density. The absorbance was normalized by the absorption peak at 260 nm. The dotted lines indicate the absorption maximum of the fluorophore. (C) Modeling fluorophore spacing in NOVA-FISH probes bound to major satellites. The B-form duplex formed by a NOVA-FISH probe (beige) and the genomic target (black) is shown. Green nucleotides indicate the locations of modified cytosines and fluorophores are depicted as green knobs. The normal distance between neighboring fluorophores in the helix is depicted. The figure was created in Pymol v.2.5.5 (RRID:SCR_000305) (63). (D) Assay to determine the impact of fluorophore number in the probe on hybridization efficiency. NOVA-FISH probes carrying increasing numbers of fluorophores (green) are hybridized with a locus of interest and dye-labeled secondary strands (red) are used as a reference. (E) Representative images of major satellites in mouse ESCs detected with NOVA-FISH probes containing increasing numbers of fluorophores. Scale bars, 5 μm. (F) Binding efficiency is unaffected by dense labeling. The normalized intensity of dye-labeled secondary strands (red) is depicted. (G) Densely labeled probes exhibit a decrease in fluorescence. Related to (F). The normalized intensity of NOVA-FISH probes (green) is depicted. Number of cells analyzed: 0% (n = 149), 25% (n = 146), 50% (n = 172), 75% (n = 147), 100% (n = 165).

**Figure 3.**
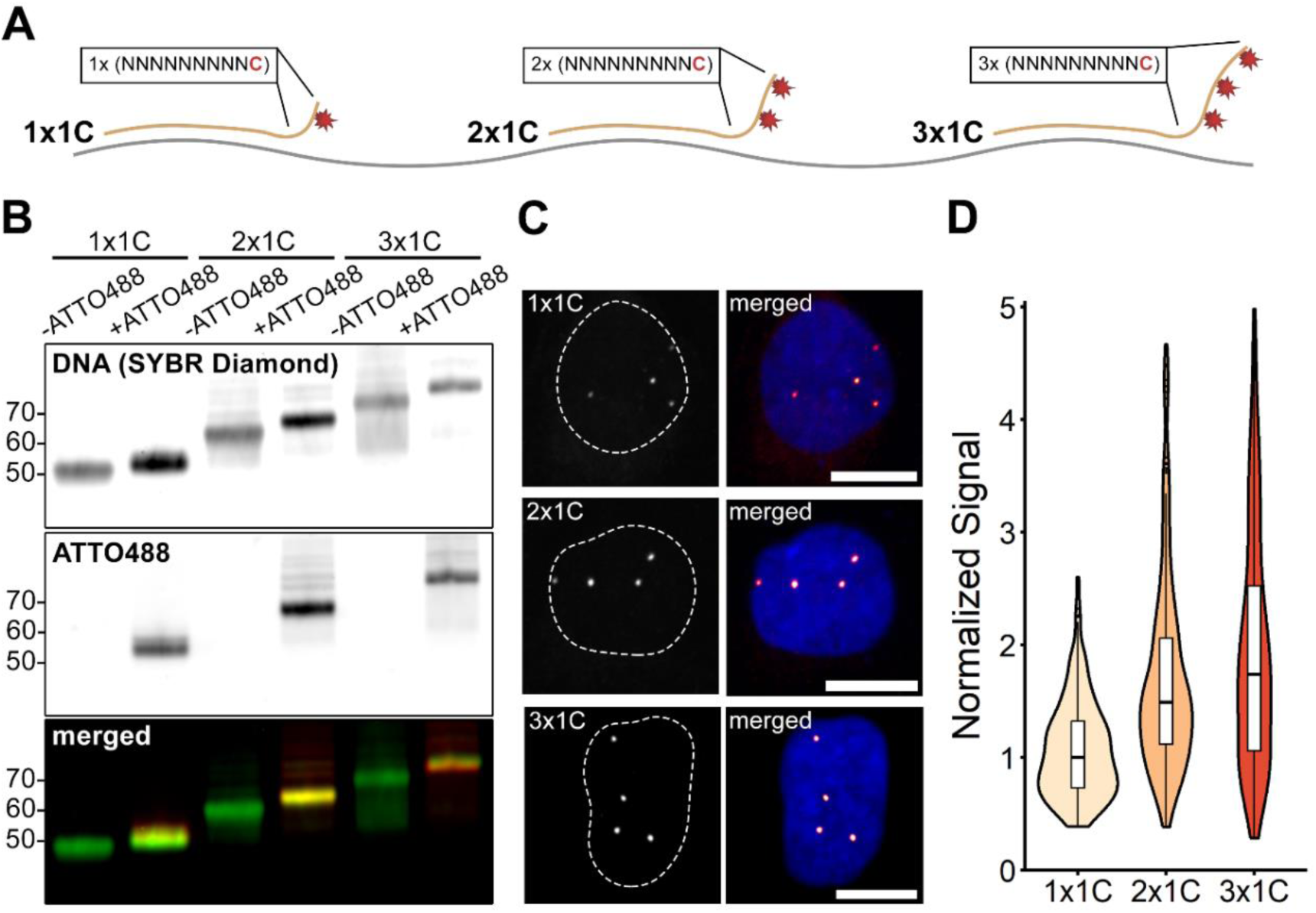
Designing extended NOVA-FISH (xNOVA) Probes. (A) Schematic of the approach. xNOVA-probes are extended by labeled 10-mers (NNNNNNNNNC) at their 3’-ends. (B) Synthesizing xNOVA-probes with specific fluorophore numbers (1x1C, 2x1C, 3x1C). Stained DNA (SYBR Diamond) and fluorophores (ATTO488) are shown in green and red, respectively. xNOVA-probes were synthesized with a 0% (-ATTO488) or 100% (+ATTO488) ratio of labeled to unlabeled nucleotide. (C) Representative images of xNOVA-probes detecting a subtelomeric region in U2OS cells. Scale bars, 10 µm. (D) Quantification of xNOVA-probe signals. Related to (C). The plot depicts the brightest signal for each cell. Number of foci analyzed: 1C (n = 270), 2x1C (n = 321), 3x1C (n = 290).

### NOVA-FISH detects non-repetitive genomic loci with kilobase resolution

Finally, we tested the limits of NOVA-FISH by detecting small non-repetitive genomic loci with nanoscale precision using STED microscopy (Figure 4A). We designed probe sets to detect non-repetitive neighboring regions on chromosome 11 termed “A” and “B” that have been established in past works ^23^. Probe sets against “A” contained 60, 50, 40, 30, 20, or 10 individual oligonucleotides, while “B” was targeted with 60 probes. The probe sets span 6.1, 4.8, 3.7, 1.7, or 0.5 kb for “A” and 4.8 kb for “B” and yield two adjacent spots (Figure 4B). A characteristic of the NOVA technology is the complete flexibility in probe synthesis as probes can be selectively amplified from a large pool by adding appropriate primer combinations (Figure 4C). This allows the cost-effective repeated use of one oligonucleotide pool to generate probes against different target regions. Then, we targeted “A” with decreasing numbers of individual probes, maintaining the same set of probes for “B” (Figure 4D). Despite observing a decline in signal intensity with the reduced number of probes detecting “A” we were still able to detect genomic loci as small as 0.5 kb. Thus, NOVA-FISH is a robust tool to detect non-repetitive regions below the kilobase level.

**Figure 4.**
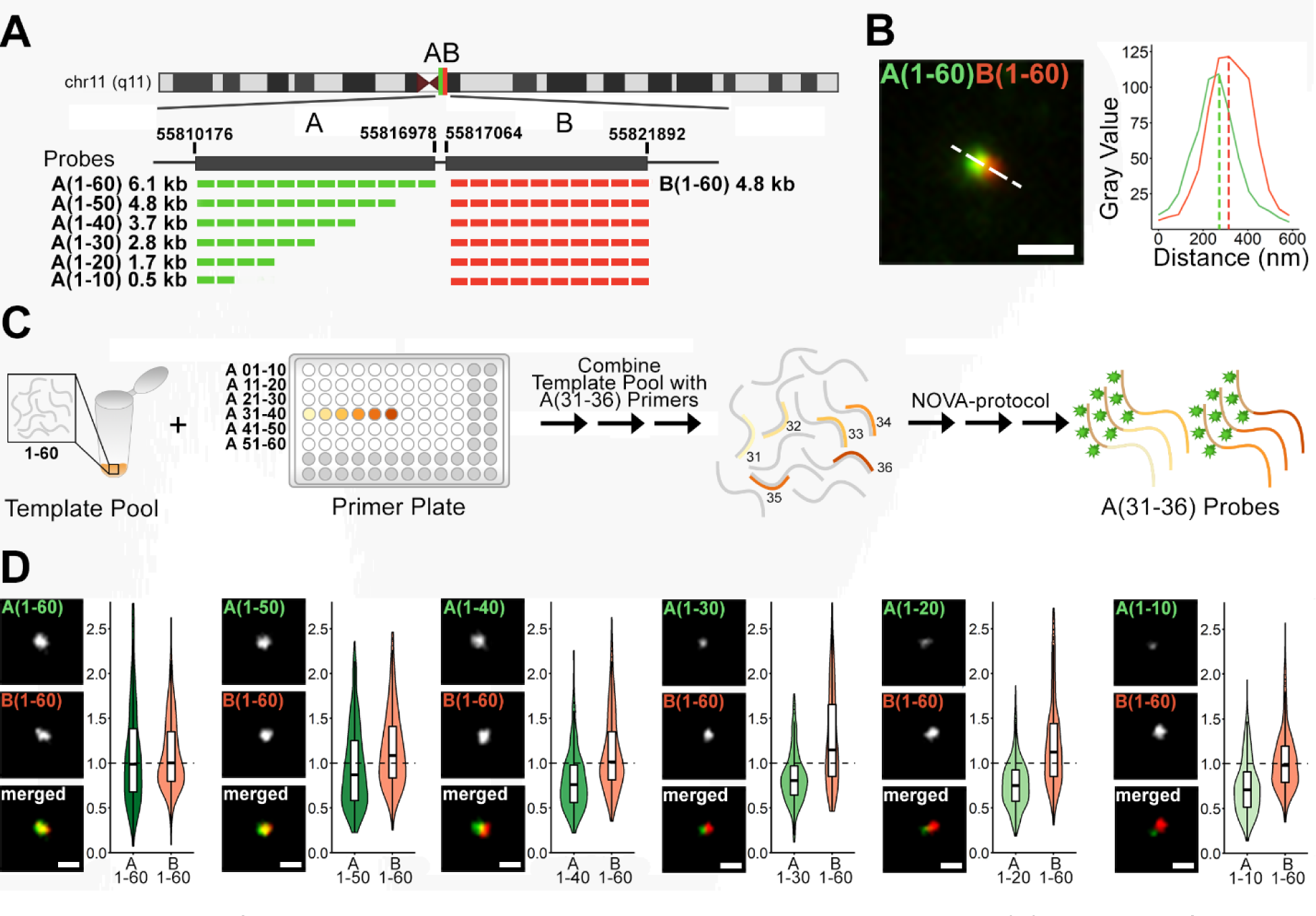
Using xNOVA-probes to detect non-repetitive loci at kilobase scale. (A) Depiction of the target regions. Two adjacent non-repetitive genomic regions on chromosome 11 (hg19, chr11: 55810176– 55816978, hg19, chr11: 55817064–55821892) were targeted with 60 xNOVA-probes spanning 6.1 or 4.8 kb. In successive experiments, the number of xNOVA-probes detecting A was gradually reduced from 60 to 10 probes. 60 xNOVA-probes targeting B were used as a reference. (B) Representative STED images of regions “A” and “B” in two colors. A(1-60) was tagged with ATTO594 and B(1-60) harbored ATTO647N. Scale bar, 500 nm. (C) Schematic of the approach. Probe sets are selectively synthesized through the combination of region-specific primers with common template pools. (D) Detecting genomic loci at kilobase resolution. 60 probes detecting B were paired with probe sets comprising 60 to 10 individual probes for A. Representative z projections of A paired with B in K562 cells are depicted. The y-axis depicts the signal intensity normalized to the median of A(1-60), B(1-60). Scale bars, 200 nm. Number of analyzed spot pairs: 1-60: n = 401, 1-50: n = 303, 1-40: n = 332, 1-30: n = 244, 1-20: n = 538, 1-10: n = 337.

## DISCUSSION

Over the past decade, it became clear that the 3D genome organization contributes to the establishment, maintenance and change of gene activity ^60,61^. Chromatin capture assays have identified genome-wide interactions of regulatory elements and have delineated TADs ^21,62^. These findings have traditionally been complemented by FISH-based imaging methods detecting entire genomes and individual chromosomes down to single genomic loci ^30,31,63,64^. However, the optical detection of small genetic elements and the resolution of their spatial relationship with promoter activity remains challenging. In this study, we developed a simple, rapid, flexible, and cost-effective protocol for the generation of FISH probe sets that are suited for nanoscopic measurements with kilobase resolution.

Small genetic elements are ideally detected with multiple synthetic oligonucleotides that may either be directly labeled or hybridized with secondary, labeled probes. Whereas end-labeled commercial probes are expensive if large and diverse probe pools are used, enzyme-based synthesis is cost-effective and flexible but requires the subsequent removal of template strands. While previously, RNA templates were reverse transcribed and subsequently degraded by RNases, we simply removed 5’-phosphorylated DNA templates using lambda exonuclease ^65^. This enzymatic synthesis, including two purification steps, takes under four hours and yields sets with hundreds of probes for less than 10 €.

For enzymatic incorporation of dye-labeled nucleotides we tested commonly available DNA polymerases. We found that B-family DNA polymerases incorporate all used modified nucleotides more effectively than A-family DNA polymerases, such as Klenow fragment or Taq DNA polymerase. This is consistent with previous structural data of B-family polymerases, attributing their ability to incorporate dye-labeled nucleotides to a larger channel volume, the presence of B-form DNA, and phosphate backbone-mediated protein-DNA interactions ^58,66–68^. Among the tested B-family polymerases, Therminator DNA polymerase having mutations in its exonuclease domain (D141A, E143A) and finger domain (A485L) was best suited for the incorporation of dye-labeled nucleotides ^49^.

As even minor FISH projects involve dozens of probes with different dye-labels, we used inexpensive, commercially synthesized template pools in combination with plates of bioinformatically optimized, target-specific primers. This approach allows the flexible generation of small to large probe sets coupled with variable dyes. We demonstrate that regions as small as 500 base pairs can be detected and genomic distances of a few kilobases can be measured.

We found that the brightness of probes can be easily adjusted with the ratio of labeled to unlabeled nucleotides in the synthesis reaction. However, the brightness did not linearly increase, due to distance-dependent effects at high labeling densities. To become independent of probe-specific sequences and to ensure incorporation of same numbers of fluorescent nucleotides we generated extended probes with overhanging, identical sequences (xNOVA). We successfully incorporated fluorophores with a spacing of ten nucleotides but assume that based on a previous study ^69^ also distances down to six nucleotides might be permissible.

While current FISH techniques can sequentially label multiple targets, the use of end-labeled probes for secondary hybridization steps reduces signal strength. To enhance signal strength, NOVA-probes carrying multiple fluorophores could be employed for secondary hybridization. Moreover, we hypothesize that our workflow is suitable for applications beyond the detection of small genomic loci. Given that oligonucleotides carrying any number of desired fluorophores can be generated, novel opportunities in the fields of DNA-PAINT, DNA origami, or immunostainings emerge ^70,71^. In summary, we present a simple, quick and inexpensive approach to explore the spatial relationships of genetic elements governing the activity of clusters of genes.

## Method Details

### Cell Culture

K562 cells were cultivated in Dulbecco’s modified Eagle’s medium (DMEM), 10% fetal bovine serum (FBS), 100 U/mL penicillin and 100 μg/mL streptomycin. U2OS cells were maintained in McCoy’s 5A medium supplemented with 10% FBS, 100 U/ml penicillin, and 100 µg/ml streptomycin at 37 °C in 5% CO_2_. IMR-90 cells were cultured in DMEM, 20% FBS, 1× MEM Non-essential amino acids, and 100 U/mL penicillin, 100 μg/mL streptomycin.

Mouse embryonic stem cells (ESCs) were maintained on culture dishes treated with 0.2% gelatin in DMEM containing 16% FBS, 0.1 mM ß-mercaptoethanol, 2 mM L-glutamine, 1× MEM Non-essential amino acids, 100 U/mL penicillin, 100 μg/mL streptomycin, homemade recombinant LIF, and 2i (1 μM PD032591 and 3 μM CHIR99021). For imaging, ESCs were seeded on plates that have been pre-treated with Geltrex diluted 1:100 in DMEM/F12 overnight at 37 °C in 5% CO_2_. Cells were passaged every 2-4 days. All cell lines were regularly tested for Mycoplasma contamination by PCR.

### Probe Design

All generated probe sets are listed in Supplementary Table S1. NOVA-probes labeling murine major satellites and human subtelomeric and telomeric regions were adapted from previously published sequences ^72,73^. Target regions (“A”: chr11:55810891-55816978 “B”: chr11:55817064-55821430) were chosen in hg38 and 60 unique oligonucleotides were selected and filtered, respectively ^23^. Barcodes of xNOVA-probes containing repetitive sequences (10-mers) were obtained from previous published data ^30^. To generate non-repetitive barcodes, pairs of orthogonal sequences from ^74^ were merged. Then, the barcodes were filtered for those containing cytosines every 10 bases and trimmed to the required length.

### NOVA-FISH Probe Synthesis

5’-phosphorylated templates and unlabeled primers were ordered from IDT or Eurofins. Equimolar amounts of 5’-phosphorylated templates and primers (0.10-0.17 nmol each) were combined to a final concentration of 1 µg DNA / µl in 1x ThermoPol® Reaction Buffer.

The annealing temperatures were adjusted to the length of the primers. For NOVA-probes (40 nt long templates, 20 nt long primers), the sample was heated up to 95 °C for 5 minutes followed by a stepwise cool-down (1 °C / minute) to room temperature. For xNOVA-probes or xNOVA-pools (50-70 nt long templates, 40 nt long primers) the sample was heated up to 95 °C for 5 minutes followed by a stepwise cool-down (1 °C / 2 minutes) to 60 °C. Complex xNOVA-probe sets were synthesized by adding two-fold excess of primer sets (e.g. primer 31-40 against “A”) to the template pool.

NOVA- and xNOVA-probes were synthesized by adding 2-4 µg annealed DNA (2-4 µl of the solution) to a reaction mixture containing 0.25 mM dATP/dGTP/dTTP each, 0-0.25 mM dCTP, 0-0.25 mM dye-labeled dCTP and 3 U Therminator DNA polymerase in 1x ThermoPol® Reaction Buffer (10 µl total volume). The ratios of dye-labeled dCTP to unlabeled dCTP varied depending on the desired labeling density. The reaction was carried out for 60 minutes at 72 °C.

To remove single-stranded DNA, NucleoSpin Gel and PCR Clean-up Kit (Macherey-Nagel) was used according to the manufacturer’s instructions. 9 volumes of buffer NTI (provided by the manufacturer) were added to one volume of sample before binding. After washing, the DNA was eluted twice in 22 µl ddH_2_O (44 µl final volume). In the next step, 5’-phosphorylated strands were removed by adding 1 µl Lambda exonuclease (10 U / µl) and 5 µl Lambda exonuclease reaction buffer (10x) to a final volume of 50 µl and incubating for 30 minutes at 37 °C. The synthesized probes were then purified using the Monarch® PCR & DNA Cleanup Kit (New England BioLabs) according to the manufacturer’s instructions and the quality was verified on denaturing 12-16% polyacrylamide gels.

### Quality Control and Purification

The absorbance of samples was measured at 260 nm and 488 nm, 596 nm, or 647 nm depending on the incorporated fluorophore using a Nanodrop™ 2000 spectrophotometer (Thermo Fisher Scientific). To assess the quality of generated probes, samples were denatured in 90% formamide, 0.5% EDTA, 0.1% Xylene cyanol, 0.1% bromophenol blue and loaded onto a 12 % polyacrylamide gel containing 6 M urea. The gel was incubated in 1x TBE buffer containing 1x Diamond™ Nucleic Acid Dye for 30 minutes at room temperature to visualize single-stranded DNA.

Complex probe sets labeling target region “A” or “B” were further purified following the “crush and soak” method with adaptations ^75^. Briefly, segments of the polyacrylamide gel containing the band of interest were cut out and 2 volumes of a buffer containing 10 mM magnesium acetate tetrahydrate, 0.5 M ammonium acetate, 1 mM EDTA (pH 8.0) and 0.1% (w/v) SDS was added followed by incubation at 37 °C for 16-24 hours. The samples were then centrifuged at 13000 x g for 1 minute and the supernatant was once more purified using the Monarch® PCR & DNA Cleanup Kit (New England BioLabs). We expect the “crush and soak” method to improve signal strength if low labeling densities are used during extension.

### Polymerase Screens

Polymerases were tested for their ability to incorporate dCTP-ATTO488, dCTP-ATTO594, or dCTP-ATTO647N into oligonucleotides. The maximum number of incorporated dCTP-dye in the probe was eight (CATCCTGAAGGAATGGTCCATG**C**TTA**CC**TGGG**CCC**AT**CC**T).

For detailed information about the reaction conditions see Supplementary Table S2. 0.1 nmol annealed DNA was added to the recommended reaction mixtures (10 µl final volume,) and 5 U of the respective polymerase was added. The following temperatures were used during synthesis: Klenow exo-at 30 °C, Taq at 64 °C, Q5 at 64 °C, Phusion at 64 °C, Therminator DNA polymerase at 72 °C. All reactions were carried out for 60 minutes and the reactions were stopped by adding 1 µl 0.5 M EDTA. We did not observe notable differences in incorporation efficiency between the reported results and reactions carried out at higher temperatures (Klenow exo-at 37 °C, Taq at 72 °C, Q5 at 72 °C, Phusion at 72 °C, Therminator DNA Polymerase at 75 °C) (figure not shown). The absorbance of synthesized products was measured at 200-700 nm on Nanoquant plates using a Tecan Spark microplate reader (Tecan) and choosing the following dye-correction factors: CF_260_(ATTO488): 0.22, CF_260_(ATTO594): 0.22, CF_260_(ATTO647N): 0.04. The depicted data contained at least two measurements per biological replicate.

### HPLC

ATTO594-labeled and ATTO647N-labeled probes (0.31 nmol or 0.34 nmol) were analyzed and purified by reverse-phase HPLC using an Agilent Technologies 1260 Infinity II System with a G7165A detector equipped with a EC 250/4 Nucleodur 100-3 C18ec column from Macherey Nagel. A gradient of 0-80% of buffer B in 45 min at 60 °C with a flow rate of 1 mL/min was applied. Following buffer system was used: buffer A: 100 mM NEt_3_ /HOAc, pH 7.0 in H_2_O and buffer B: 100 mM NEt_3_ /HOAc, pH 7.0 in H_2_O/MeCN 20/80. The fractions of each signal peak were combined, and the solvents were concentrated by vacuum centrifugation.

### Sample preparation and fluorescence in situ hybridization

Fluorescence in situ hybridization was performed as previously described ^23^. Adherent cells were grown overnight on glass coverslips (1.5, 18x18 mm, Marienfeld), washed twice with 1x Dulbecco’s Phosphate Buffered Saline (PBS), and fixed using osmotically balanced and methanol-free 4% formaldehyde for 10 minutes at room temperature. Alternatively, PBS-washed suspension cells were resuspended in a small volume of PBS at a density of 1 million cells per ml and applied to poly-L-lysine coated glass coverslips followed by the addition of methanol-free 4% formaldehyde for 10 minutes at room temperature. The slides were washed twice in 1x PBS for 5 minutes and the cells were permeabilized in 1X PBS containing 0.5% Triton X-100 for 15 minutes. After two successive washing steps in 1x PBS, 0.1 M HCl was added to the slides for 5 minutes. The slides were washed twice with 2 x SSC and were placed onto a solution containing 1 µg/ml RNase for 30 minutes at 37 °C in a wet chamber. Then, adherent or suspension cells were pre-equilibrated in 2x SSC containing 50% formamide for 60 minutes or overnight, respectively, inverted onto 8 µl of hybridization solution, and sealed with rubber cement (Marabu). The slides were placed on a heat block set to 81 °C for 3 minutes and incubated at 37 °C overnight (16 – 20 hours).

On the second day, slides were washed twice with 2x SSC for 15 minutes followed by two successive 7-minute washes in 0.2x SSC containing 0.2% Tween-20 at 56 °C. Then, slides were washed with 4x SSC containing 0.2% Tween20 and with 2x SSC for 5 minutes, respectively.

For oligoFISH probes, a second hybridization step was performed for 30 minutes at room temperature. The slides were then washed once with 2x SSC containing 30% formamide for 7 minutes at 37 °C, twice with 2 x SSC for 5 minutes, once with 0.2X SSC containing 0.2% Tween20 at 56 °C, once with 4x SSC containing 0.2% Tween20 for 7 minutes at room temperature and once with 2x SSC for five minutes.

DNA was counterstained with DAPI (1 µg/ml in 2x SSC) for 10 minutes and washed twice with 2x SSC. For STED microscopy, nuclei were counterstained with or DiYO-1 (12.5 nM in 2x SSC) for 30 minutes and washed twice with 2x SSC for 5 minutes, respectively. Coverslips were mounted on microscopic slides with MOWIOL (2.5% DABCO, pH 7.0), dried for 30 minutes, and sealed with nail polish.

### Statistics and reproducibility

The experiments shown in this study were performed as three biologically independent experiments (n = 3) and the figures contain pooled data. No statistical methods were used to predetermine the sample size. Images depicted are representative images from the experiments and dotted lines indicate the outlines of the cells. Data plotted as boxplots indicate the 25th and 75th percentiles, with the whiskers showing the minima and maxima (5th and 95th percentiles), black circles indicating the outliers, and the horizontal line showing the median. Some data are plotted in bar graphs as the mean ± s.d. Data was normalized by the median of the first depicted condition in the replicates, if not stated otherwise. Significance levels were tested by non-parametric two-sided Wilcoxon tests or pairwise comparisons using the Wilcoxon rank sum test with Bonferroni’s correction for multiple testing (* = p < 0.05, ** = p < 0.01, *** = p < 0.001). Sample sizes for all of the graphs are indicated in the figures or figure legends.

### Image Acquisition

Confocal images were acquired using a Nikon TiE microscope equipped with a Yokogawa CSU-W1 spinning-disk confocal unit (50 μm pinhole size), an Andor Borealis illumination unit, Andor ALC600 laser beam combiner (405 nm/488 nm/561 nm/640 nm), Andor IXON 888 Ultra EMCCD camera, and a Nikon 100×/1.45 NA oil immersion objective. The microscope was controlled by software from Nikon (NIS Elements, ver. 5.02.00).

Super-resolution was carried out on a 2C STED 775 QUAD Scan microscope (Abberior Instruments) equipped with a 100x 1.4 NA UPlanSApo oil immersion objective lens (Olympus), 3 pulsed excitation lasers (485 nm, 594 nm, 640 nm) and a pulsed depletion laser of 775 nm.

### 3D STED microscopy of telomers using adaptive illumination

To avoid photobleaching NOVA-FISH stained telomers of IMR90 cells in 3D, stacks were acquired using adaptive illumination STED microscopy ^76^. Cells were recorded using a pixel size of 30 nm, z-steps of 80 nm, a 10 µs dwell time, and a pinhole size of 50 µm.

### Automated STED microscopy for two-color NOVA-FISH

Automated STED microscopy was performed according to Brandstetter et al. ^23^. The acquisition of 3D confocal stacks was automated using home-written Python scripts to control the microscope. Spots within confocal scans were detected using a Laplacian-of-Gaussian blob detector for both channels. Detected spots no further apart than 5 pixels from another spot in the other channel were imaged using 3D STED settings. This process was repeated for each detected spot pair within a confocal scan. Following a spiral pattern, the stage was moved to the next overview to repeat the confocal scan and the subsequent detailed STED acquisition until a specified amount of time elapsed.

### Image Analysis

For the analysis of the effects of labeling density (Figure 2E-G), cells in confocal z-stacks of major satellites were segmented first via automatic thresholding in a z-maximum projection of the DAPI channel followed by a second round of thresholding in the 640nm (rel. binding) channel to segment major satellites. In the segmented areas, intensities of both the 488nm (rel. brightness) and the 640nm (rel. binding) channels were measured, background determined by a manually selected ROI outside the cells was subtracted, and measurements were averaged (median) per cell. For the plots, measurements were normalized to the intensity at 100% for the binding channel and at 25% for the brightness channel. Analysis was carried out using Fiji ^77^.

For analysis of image data of telomeres and subtelomeric regions (Figure 1B-C, Figure 3C-D, Figure S7A-B), nuclear segmentation maps of confocal images stained with DAPI or YOYO-1 were obtained using Otsu thresholding. FISH spots within segmentation maps were detected using a Laplacian-of-Gaussian blob detector. Segmentation maps were used to calculate the total number of spots per cell, to obtain the mean background signal within single nuclei to calculate the spot signal over the nucleus background, and the signal-to-noise ratio of single spots.

Analysis of automated STED measurements of FISH spot pairs was performed as previously described ^23^. Automated image acquisition generated large quantities of data requiring an additional quality control step. To filter out low-quality images, we used a machine learning-based classifier (Random Forest) to label images as “good” or “bad”. The classifier was trained with a ground truth dataset created by an experienced scientist who manually sorted images.

Detailed spot analysis was performed on images passing this QC step. Subpixel localization of FISH spots in both channels was performed by fitting a multidimensional Gaussian function plus a constant background using the Levenberg-Marquardt algorithm. The peak height of the fitted Gaussians was used to determine spot intensity.

## ACKNOWLEDGMENTS

We thank Cristina Cardoso and Irina Solovei for helpful discussions and input. This work was supported by grants from the Deutsche Forschungsgemeinschaft [SFB1064 / project number 213249687 to HL and Priority Program SPP 2202 / project number 422857584 to HH and HL] and the BMBF Clusters4Future program CNATM. CS is a fellow of the International Max Planck Research School for Molecular Life Sciences (IMPRS-LS). Microscopic images were acquired at microscopes of the Center for Advanced Light Microscopy (CALM) at LMU Munich.

## AUTHOR CONTRIBUTIONS

C.S, H.L and H.H designed the study and wrote the manuscript. H.L., H.H., T.C. supervised the study. C.S., M.G.O, G.S., D.H., and A.J.T. performed experiments and prepared Figures. All authors reviewed the manuscript.

---

**Supplementary Figure 1.**
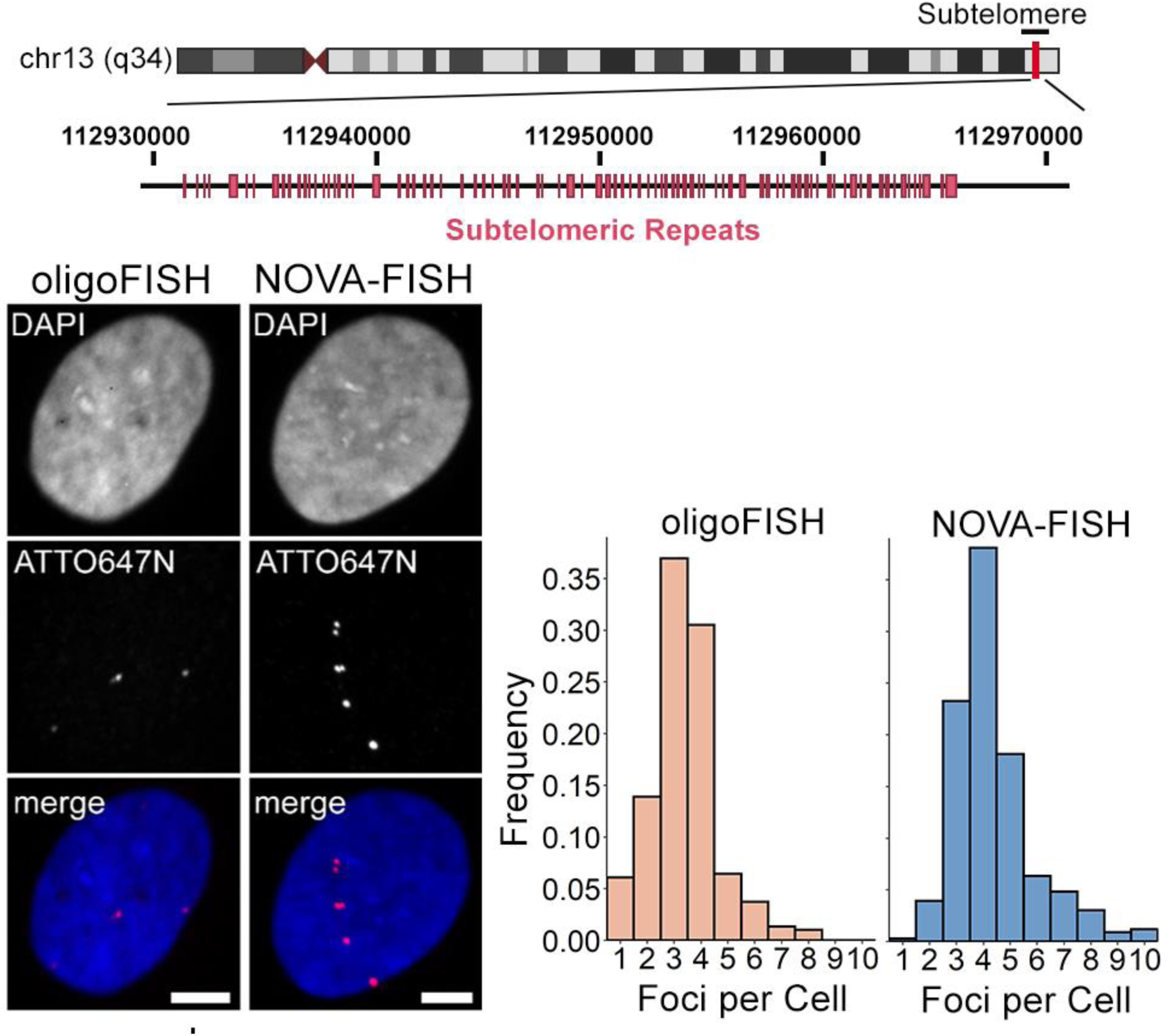
NOVA-probes facilitate the detection of polyploidies. Top: Detecting a subtelomeric region on human chromosome 13. The target region is located in chr13 q34 (hg19, chr13:112931779 112966593) and contains a unique series of repeats (pink). Bottom: Representative images for oligoFISH and NOVA-FISH detecting the subtelomeric region in U2OS cells. Scale bars, 5 μm. Histogram of the number of foci detected in each cell. Average number of detected loci: oligoFISH = 3.39, NOVA-FISH = 4.39.

**Supplementary Figure 2.**
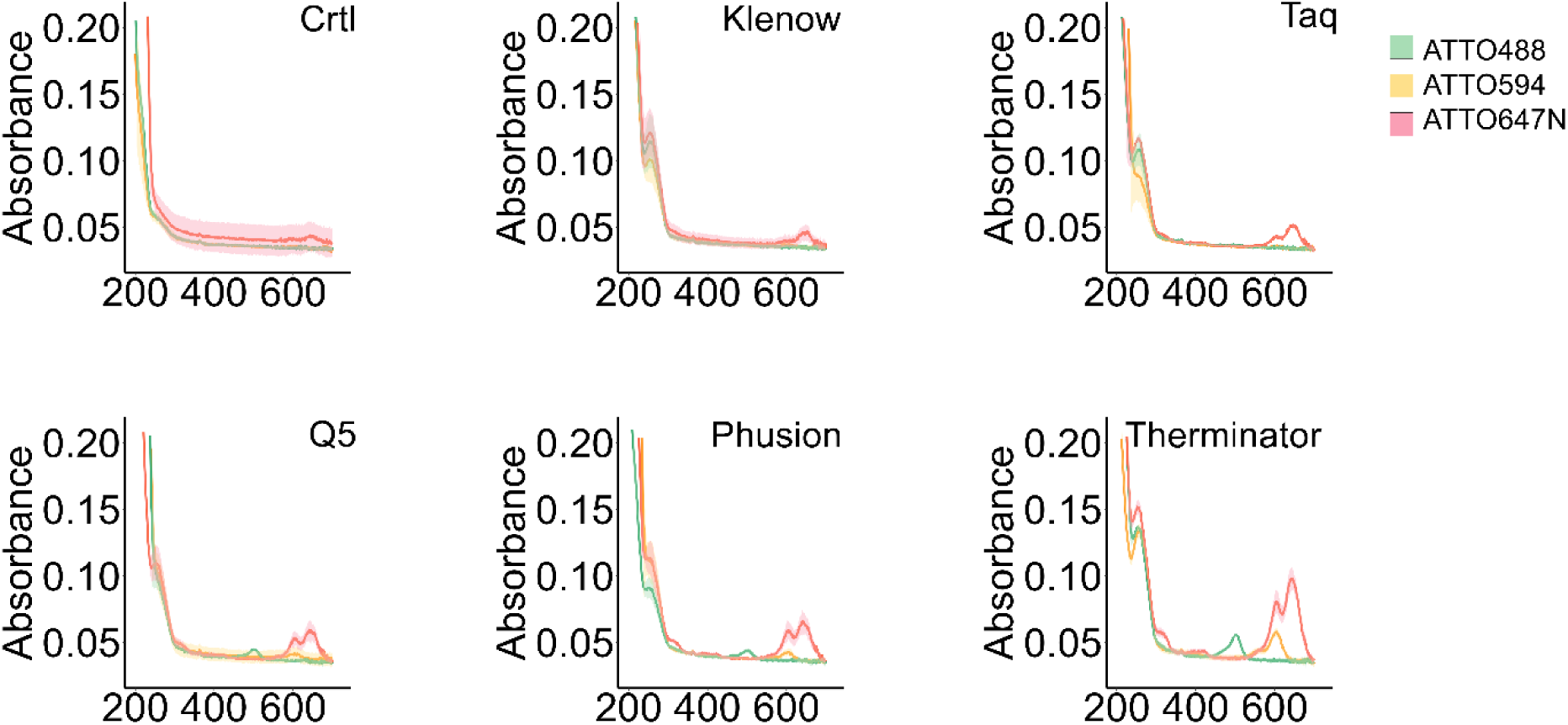
Absorption spectra of synthesized probes. The line depicts the mean absorption and the area indicates the standard deviation of three experiments with at least two separate measurements (n = 3). The incorporated nucleotides are indicated as followed: ATTO488 = green, ATTO594 = yellow, ATTO647N = red.

**Supplementary Figure 3.**
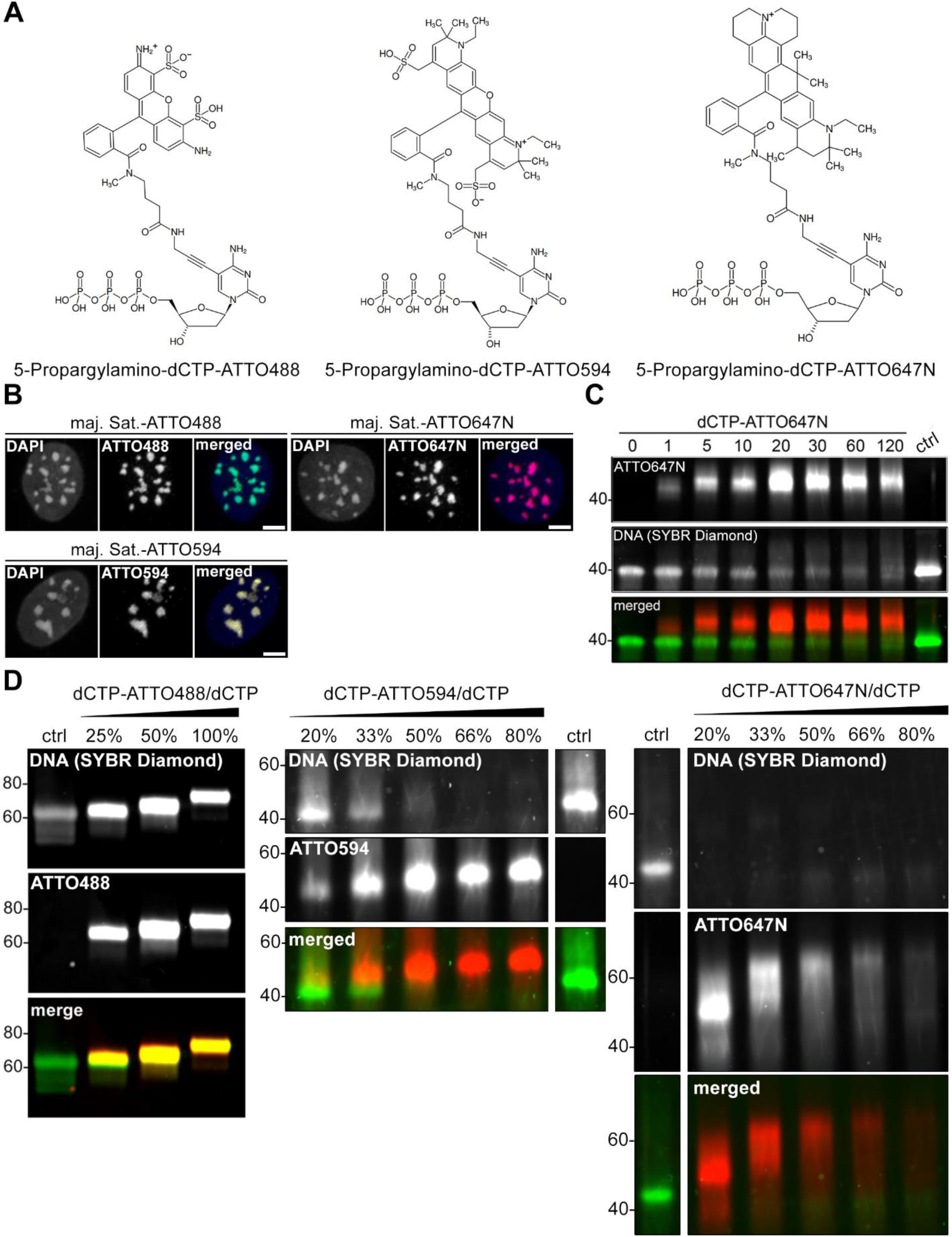
Synthesizing NOVA-probes harboring different fluorophores. (A) Structural formulas of used modified nucleotides. Dyes (ATTO488, ATTO594, ATTO647N) are linked to 5C positions of cytosines. The structures were provided by the manufacturer (Jena Bioscience). (B) Labeling major satellites in J1 cells with NOVA-FISH using three different fluorophores. All probes (maj. Sat.-ATTO488, maj. Sat.-ATTO594, maj. Sat.-ATTO647N) were generated using a one to four molar ratio of dye-labeled to unlabeled nucleotides. Scale bars, 5 µm (C) Therminator DNA polymerase effectively generates dye-labeled probes. The synthesis was carried out between 0-120 minutes using 0.15 nmol DNA and 3 U Therminator polymerase. Fluorescent DNA (ATO647N) and DNA (SYBR-Diamond) are shown in red and green, respectively. (D) Generating probes with different labeling densities. Left: Synthesis of ATTO488-labeled probes detecting major satellites. Middle: Synthesis of ATTO594-labeled probes targeting a subtelomeric region in chromosome 13. Right: Synthesis of ATTO647N-labeled probes detecting a subtelomeric region in chromosome 13. Different dye-labeled dCTP to dCTP ratios were used (25%, 50%, 100% or 20%, 33%, 50%, 66% 80%). Unlabeled probes were used as a control. Fluorophores and stained DNA (SYBR Diamond) are shown in red and green, respectively.

**Supplementary Figure 4.**
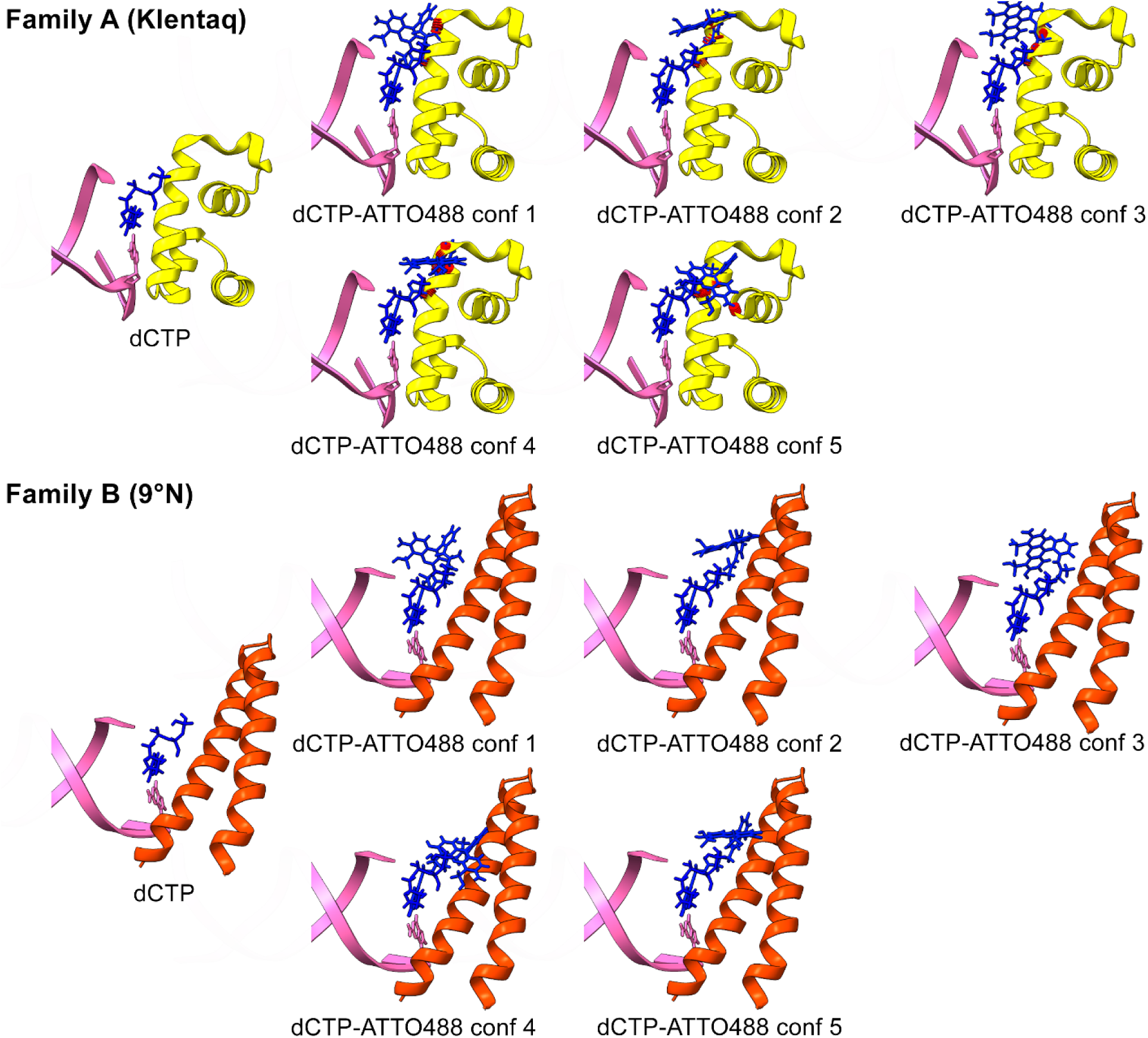
Modeling the proximity of dCTP-ATTO488 to family A or family B finger domains. Finger domains are shown in yellow (Klentaq) and orange (9°N DNA polymerase) with DNA in pink and the substrate in blue. Five different conformations of dCTP-ATTO488 (conf1-5) were superimposed on cytosine. Distances between the finger domain and dCTP-ATTO488 < 1 Å are depicted as red knobs. The finger domains are shown in the closed state. The figure was generated with UCSF Chimera (v.1.17.3, RRID:SCR_015872) by using the structures 3RTV and 5OMV ^58,59,78^.

**Supplementary Figure 5.**
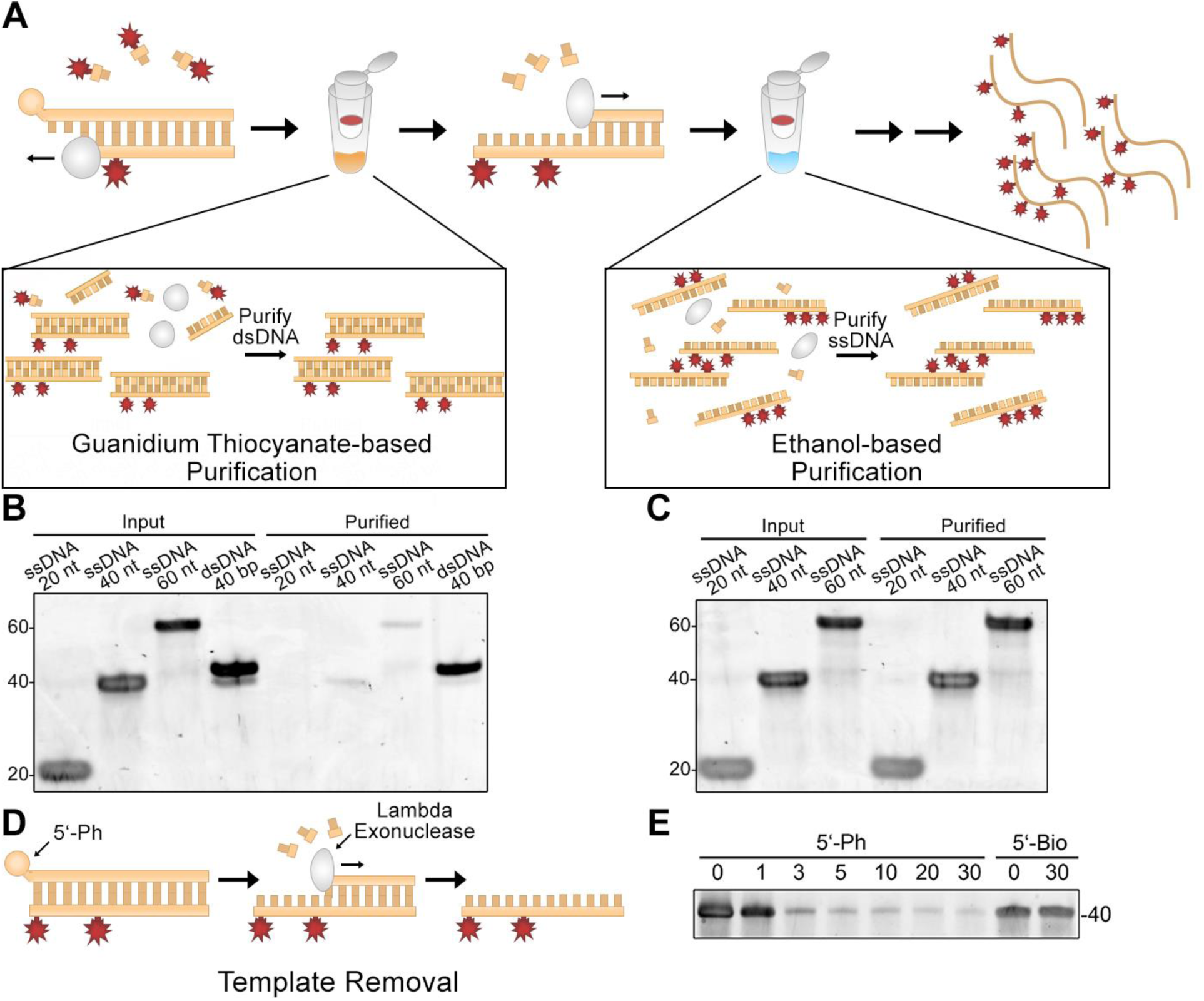
Optimization of probe purification. (A) Schematic highlighting two purification steps in the protocol. While guanidium thiocyanate-based purification enriches double-stranded DNA, ethanol-based purification purifies single-stranded oligonucleotides. (B) Guanidium thiocyanate-based purification yields double-stranded DNA. ssDNA (20 nt, 40 nt, 60 nt) and dsDNA (40 bp) were loaded before (input) and after (purified) purification. (C) Ethanol-based purification yields single-stranded DNA. ssDNA (20 nt, 40 nt, 60 nt) was loaded before (input) and after (purified) purification. (D) Schematic of lambda exonuclease-mediated degradation. Lambda exonuclease selectively removes 5’-phosphorylated (5’-Ph) strands. (E) Removal of the 5’-phosphoryated template. Templates were incubated with 10 U lambda exonuclease. 5-biotinylated oligonucleotides were used as a control.

**Supplementary Figure 6.**
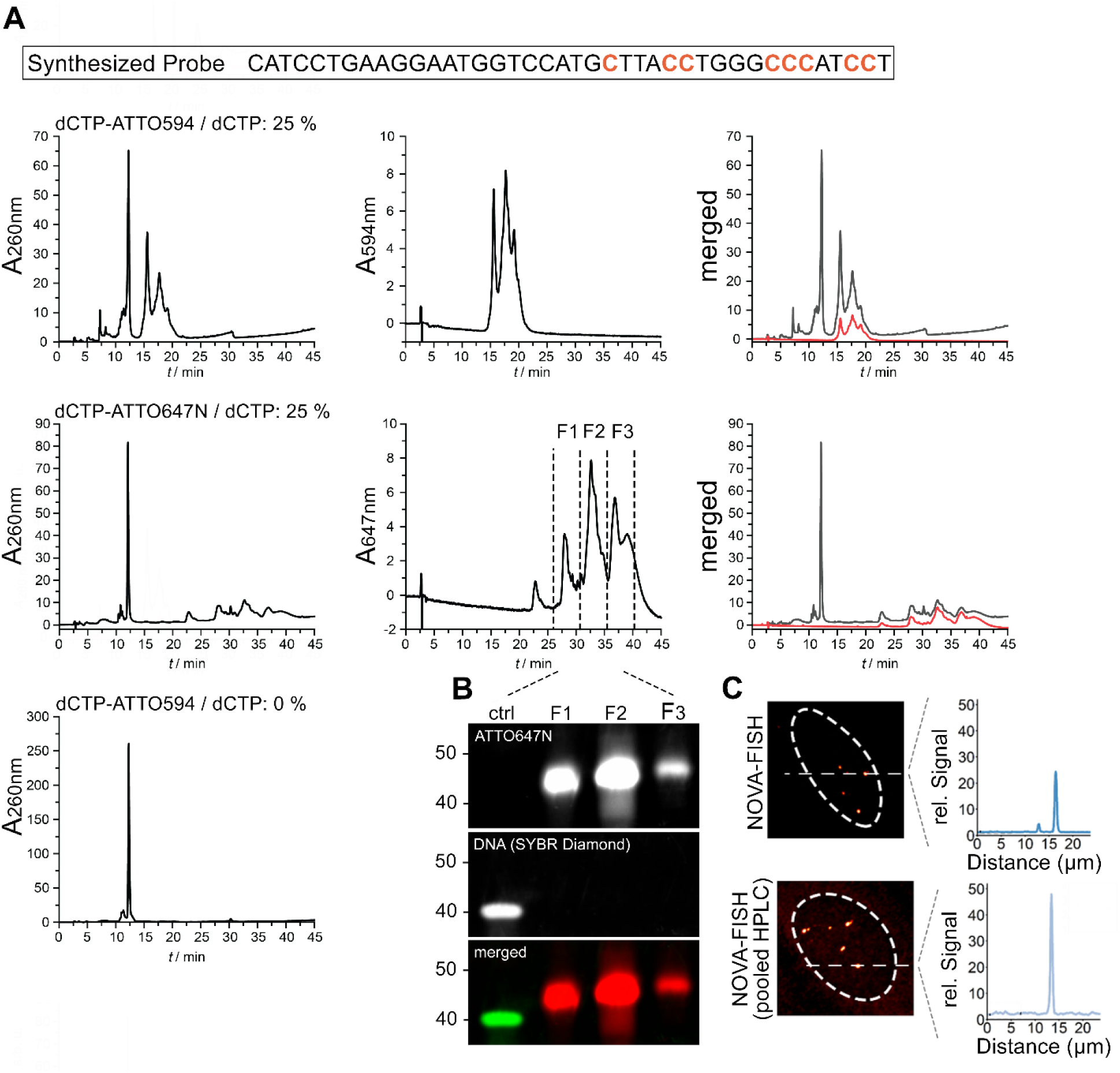
Characterizing NOVA-probes with low fluorophore input in synthesis. (A) HPLC of NOVA-FISH probes. Top: Sequence of the synthesized probe. Possible incorporation sites of modified nucleotides are highlighted in red. Middle: Elution profile of synthesized probes. Probes were synthesized with Therminator DNA Polymerase using a one to four molar ratio of dye-labeled to unlabeled nucleotides. The absorptions of ATTO594-labeled and ATTO647N-labeled probes were measured at 260 nm / 594 nm and 260 nm / 647 nm, respectively. Local peaks indicate probes with varying numbers of fluorophores. Probes synthesized without modified nucleotides were used as a control (bottom left). (B) Polyacrylamide gel reveals a visible shift between populations with different fluorophore numbers. Three fractions (F1, F2, F3) were isolated from (A) through preparative HPLC. The precise mechanism of SYBR-Diamond-ssDNA interactions has yet to be elucidated but we observed that the presence of fluorophores in the probe impacts staining efficiency. Unlabeled DNA was used as a control (ctrl). (C) Side-by-side comparison of unpurified and purified NOVA-probes. To evaluate the benefits of HPLC purification, we conducted FISH in U2OS cells using NOVA-probes before and after HPLC purification. The relative signal intensity along the dotted line is depicted. Images were acquired at the same conditions.

**Supplementary Figure 7.**
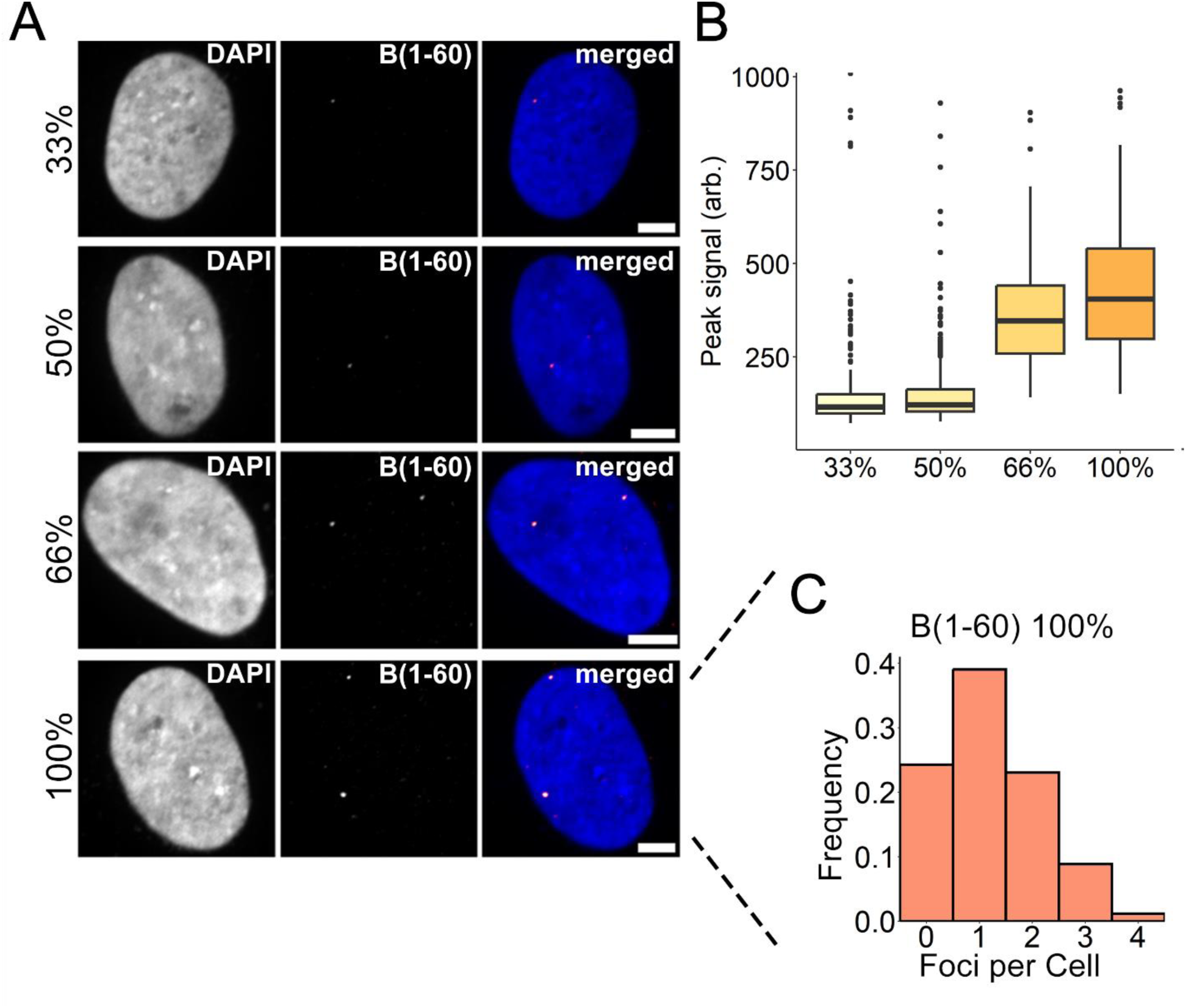
Increased labeling densities in xNOVA-probes. (A) Representative confocal image of a non-repetitive region (“B”, see Figure 4A) in U2OS cells. “B” was labeled with 60 xNOVA-probes containing increasing labeling densities (33%, 50%, 66%, 100%). Scale bars, 5 μm. (B) xNOVA-FISH signals increase with higher labeling densities. Detected FISH intensities are displayed as boxplots. The signal was normalized by the background. (C) Histogram of the number of foci detected in U2OS cells (n = 169). Related to (A). The values were normalized by the total number of cells.

